# SARS-CoV-2 accessory proteins involvement in inflammatory and profibrotic processes through IL11 signaling

**DOI:** 10.1101/2023.03.27.534381

**Authors:** Blanca Dies López-Ayllón, Ana de Lucas-Rius, Laura Mendoza-García, Tránsito García-García, Raúl Fernández-Rodríguez, José M. Suárez-Cárdenas, Fátima Milhano Santos, Fernando Corrales, Natalia Redondo, Federica Pedrucci, Sara Zaldívar-López, Ángeles Jiménez-Marín, Juan J. Garrido, María Montoya

## Abstract

SARS-CoV-2, the cause of the COVID19 pandemic, possesses eleven accessory proteins encoded in its genome. Their roles during infection are still not completely understood. Transcriptomic analysis revealed that both *WNT5A* and *IL11* were significantly up-regulated in A549 cells expressing individual accessory proteins ORF6, ORF8, ORF9b or ORF9c from SARS-CoV-2 (Wuhan-Hu-1 isolate). IL11 signaling-related genes were also differentially expressed. Bioinformatics analysis disclosed that both *WNT5A* and *IL11* were involved in pulmonary fibrosis idiopathic disease. Functional assays confirmed their association with profibrotic cell responses. Subsequently, data comparison with lung cell lines infected with SARS-CoV-2 or lung biopsies from patients with COVID19 evidenced altered gene expression that matched those obtained in this study. Our results show ORF6, ORF8, ORF9b and ORF9c involvement in inflammatory and profibrotic responses. Thus, these accessory proteins could be targeted by new therapies against COVID19 disease.

**Research topic(s):** Viral diseases, COVID19 insights

## Introduction

The coronavirus disease 2019 (COVID19) is a potentially fatal respiratory disease caused by the new Severe Acute Respiratory Syndrome Coronavirus 2 (SARS-CoV-2), which rapidly spread worldwide causing more than 670 million reported cases and nearly 7 million deaths globally since the start of the pandemic (https://coronavirus.jhu.edu/map.html). The clinical course of COVID19 exhibits a broad spectrum of severity and progression patterns. While the infection leads to mild upper respiratory disease or even asymptomatic sub-clinical infection in a significant number of people, others develop symptoms and complications of severe pneumonia that can be fatal. Furthermore, pulmonary fibrosis has been described as one of the most Fabbri et al. estimated that approximately 20% of patients with COVID19 had evidence Since March 2020, many efforts have been done to elucidate COVID19 pathogenesis, but the complete clinical picture following SARS-CoV-2 infection is not yet fully understood.

Like the rest of Coronaviruses, SARS-CoV-2 genome consists of a single-stranded positive-sense RNA molecule of approximately 29,900 nucleotides (NCBI Reference Sequence: NC_045512.2) arranged into 14 open reading frames (ORFs) and encoding 31 proteins ^7^. Following a typical 5’-3’ order of appearance, SARS-CoV-2 proteins comprise two large polyproteins: ORF1a and ORF1b; four structural proteins: spike (S), envelope (E), membrane (M), and nucleocapsid (N) and eleven accessory proteins: ORF3a, ORF3b, ORF3c, ORF3d, ORF6, ORF7a, ORF7b, ORF8, ORF9b, ORF9c and ORF10 ^8–11^. As their name suggest, accessory proteins are dispensable for viral replication, but recent reports have demonstrated their involvement in COVID19 pathogenesis by mediating antiviral host responses ^12–15^.

SARS-CoV-2 ORF6 is a 61 aa protein that localizes in endoplasmic reticulum and membrane of vesicles such as autophagosomes and lysosomes ^9, 16^. This accessory protein displays multifunctional activities such as blocking nucleopore movement of newly synthetized mRNA encoding immune-modulatory cytokines such as IFN-β and interleukin-6 (IL-6) counteracting those cytokines ^17^. SARS-CoV-2 ORF8 is a 121 aa protein consisting of an N-terminal signal sequence for endoplasmic reticulum (ER) import. It is a secreted protein, rather than being retained in the ER, and its extracellular form has been detected in the supernatant of cell cultures and sera of COVID19 patients ^9, 18^. In addition, ORF8’s functions are mediated by its binding to CD16a, decreasing the capacity of monocytes to mediate antibody-dependent cellular cytotoxicity (ADCC) ^19^. SARS-CoV-2 ORF9b is a 97 aa protein that antagonizes type I and III interferons by negatively regulating antiviral immunity ^20^. It is localized in the mitochondrial membrane associated with TOM70 ^13^ inducing pro-inflammatory mitochondrial DNA release in inner membrane-derived vesicles ^21^. SARS-CoV-2 ORF9c is a 73 aa membrane-associated protein that suppresses antiviral responses in cells ^22^. It also interacts with Sigma receptors that are implicated in lipid remodeling and ER stress response ^9, 23^.

SARS-CoV-2 mostly affects the respiratory tract usually leading to pneumonia in most patients, and to acute respiratory distress syndrome (ARDS) in 15% of cases. ARDS is mainly triggered by elevated levels of pro-inflammatory cytokines, such as Interleukin 6 (IL6), referred to as cytokine storm ^24^. Interleukin 11 (IL11) is a member of the IL6 family of cytokines. IL11 is similar to IL6, and both form a GP130 heterodimer complex to initiate its downstream signaling ^25–28^, but their respective hexameric signaling complex formation differ ^29^. While IL6R is expressed most highly on immune cells, IL11RA is expressed in stromal cells, such as fibroblasts and hepatic stellate cells, and also on parenchymal cells, including hepatocytes. Hence, it may be expected that IL6 biology relates mostly to immune functions whereas IL11 activity is more closely linked to the stromal and parenchymal biology ^25, 30–32^. Since the nineties, high IL11 release during viral infections have been described ^33, 34^, and more recently, several studies have related this interleukin to fibrosis, chronic inflammation and matrix extracellular remodeling ^31, 35–39^. It is also known that WNT5A and IL11 have the ability of activating STAT3 signaling ^40^ and this ability has been postulated as a possible mechanism to link *WNT5A* gene with immunomodulation. WNT5A is a member of WNT family proteins which plays critical roles in a myriad of processes in both health and disease, such as embryonic morphogenesis, fibrosis, inflammation or cancer ^41^. Several studies have described a crosstalk between transforming growth factor-beta (TGFβ) and WNT signaling pathways during fibrotic processes ^42–45^, and more recently with the increase in IL11 production ^46^. TGFβ represents the most prominent profibrotic cytokine by upregulating production of extracellular matrix (ECM) components and multiple signaling molecules ^47^.

It is known that the underlying cause of severe COVID19 disease is a cytokine dysregulation and hyperinflammation status ^24, 48, 49^, and IL6 was from the beginning involved as it was found to be elevated in serum of COVID19 patients ^50, 51^. However, little is known about the involvement of IL11 in lung fibrosis in COVID19 disease. In this study, A549 lung epithelial cells were individually transduced with accessory proteins ORF6, ORF8, ORF9b or ORF9c from SARS-CoV-2 (Wuhan-Hu-1 isolate), and transcriptomic analysis revealed that both, *WNT5A* and *IL11*, were significantly up-regulated. IL11 signaling-related genes, such as *STAT3* or *TGF*β, were also differentially expressed. Subsequently, bioinformatics and functional assays revealed that these four accessory proteins were implicated in both inflammatory and fibrotic responses, suggesting the involvement of ORF6, ORF8, ORF9b and ORF9c in inflammatory and/or fibrotic responses in SARS-CoV-2 infection.

## Results

### Expression of SARS-CoV-2 ORF6, ORF8, ORF9b or ORF9c accessory proteins alter gene expression pattern in A549 cells

SARS-CoV-2 uses several strategies to interact and interfere with the host cellular machinery. To explore the function of individual ORFs in such interaction, A549 human lung carcinoma cells were lentivirus transduced expressing individual viral accessory proteins ORF6, ORF8, ORF9b or ORF9c (Figure 1A), with a C-terminally 2xStrep-tag to facilitate detection of their expression (named ORF-A549 thereafter). GFP-lentivirus-transduced or wild-type A549 cells were used as control in each experiment, both giving the same results. ORFs overexpression in A549 transduced cells was verified by immunofluorescence staining using anti-StrepTag antibody which highlighted different patterns of localization in A549 cells as well as variable levels of expression (Figure 1B). ORF9b and ORF9c seemed to be highly concentrated around the nucleus, while ORF6 and ORF8 were localized mainly in a specific perinuclear region.

**Figure 1.**
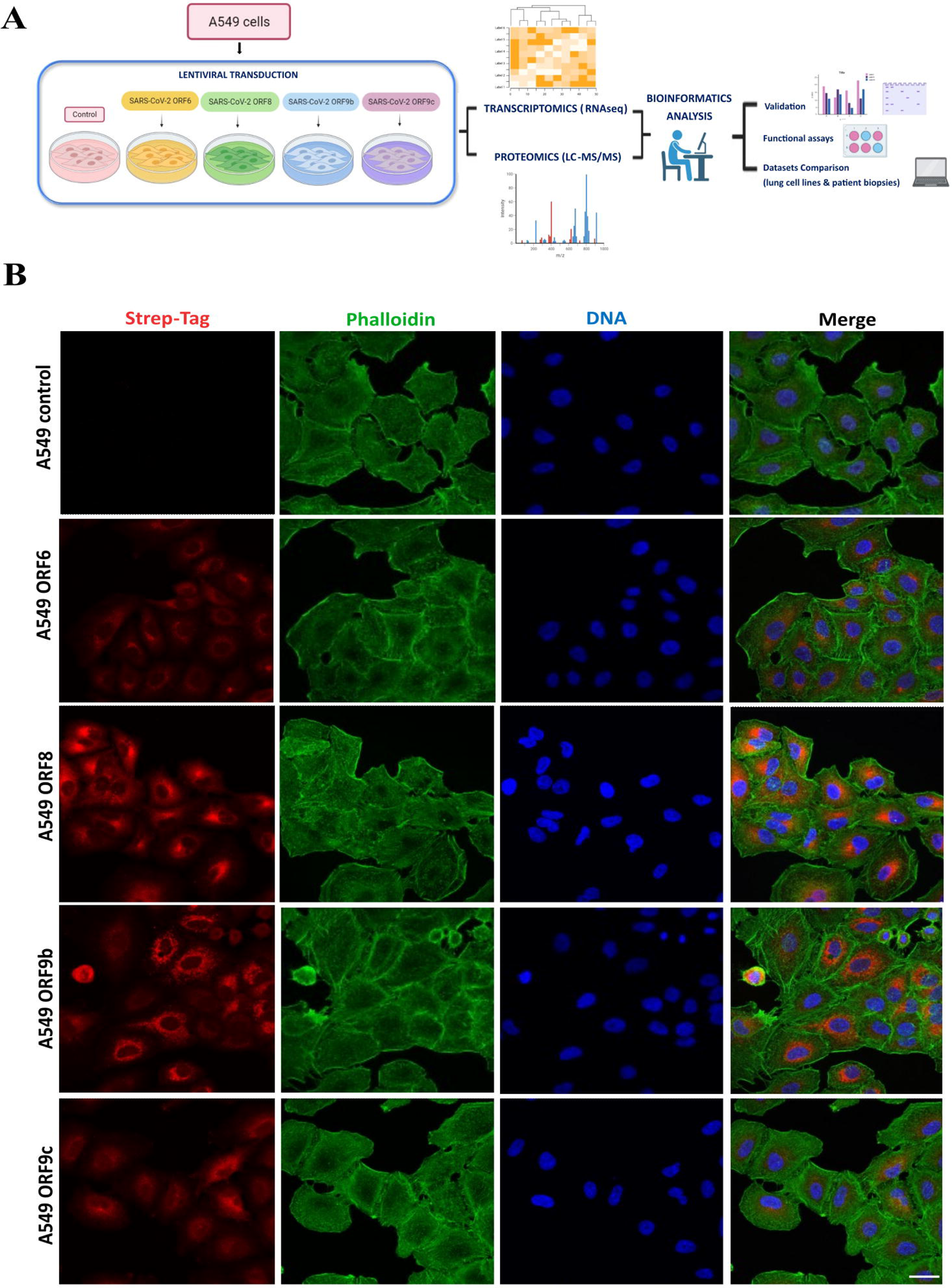
Expression of SARS-CoV-2 ORF6, ORF8, ORF9b or ORF9c in A549 epithelial cells. **A)** Experimental workflow scheme. Figure generated in Biorender. **B)** Cellular localization of ORF6, ORF8, ORF9b or ORF9c. A549 transduced cells with Strep-tagged viral proteins were imaged by confocal microscopy. Red: Strep-tag antibody signal; Green: Phalloidin; Blue: DAPI (nuclei staining). Objective 63x, scale bar 25 µm.

Differential gene expression analysis was performed in ORF-A549 cells (Figure 2A). Sample quality control was assessed by principal component analysis (PCA) based on normalized counts from DESeq2. High quality was achieved since samples were clustered (Figure 2B). Further analysis of transcriptomics data revealed a number of genes commonly expressed in all transduced cells, including *WNT5A* and *IL11*. These two genes were particularly upregulated, as well as other genes previously related to their signaling pathways ^40, 41, 52^ (Figure 2C). qRT-PCR was used to validate transcriptomic data in ORF-A549 cells for *CXCL1, IL11, WNT5A, WNT5A-AS* and *STAT3* (Figure 2D). Also, IL11 release was significantly increased in cells expressing ORF8, ORF9b and ORF9c (Figure 2E).

**Figure 2.**
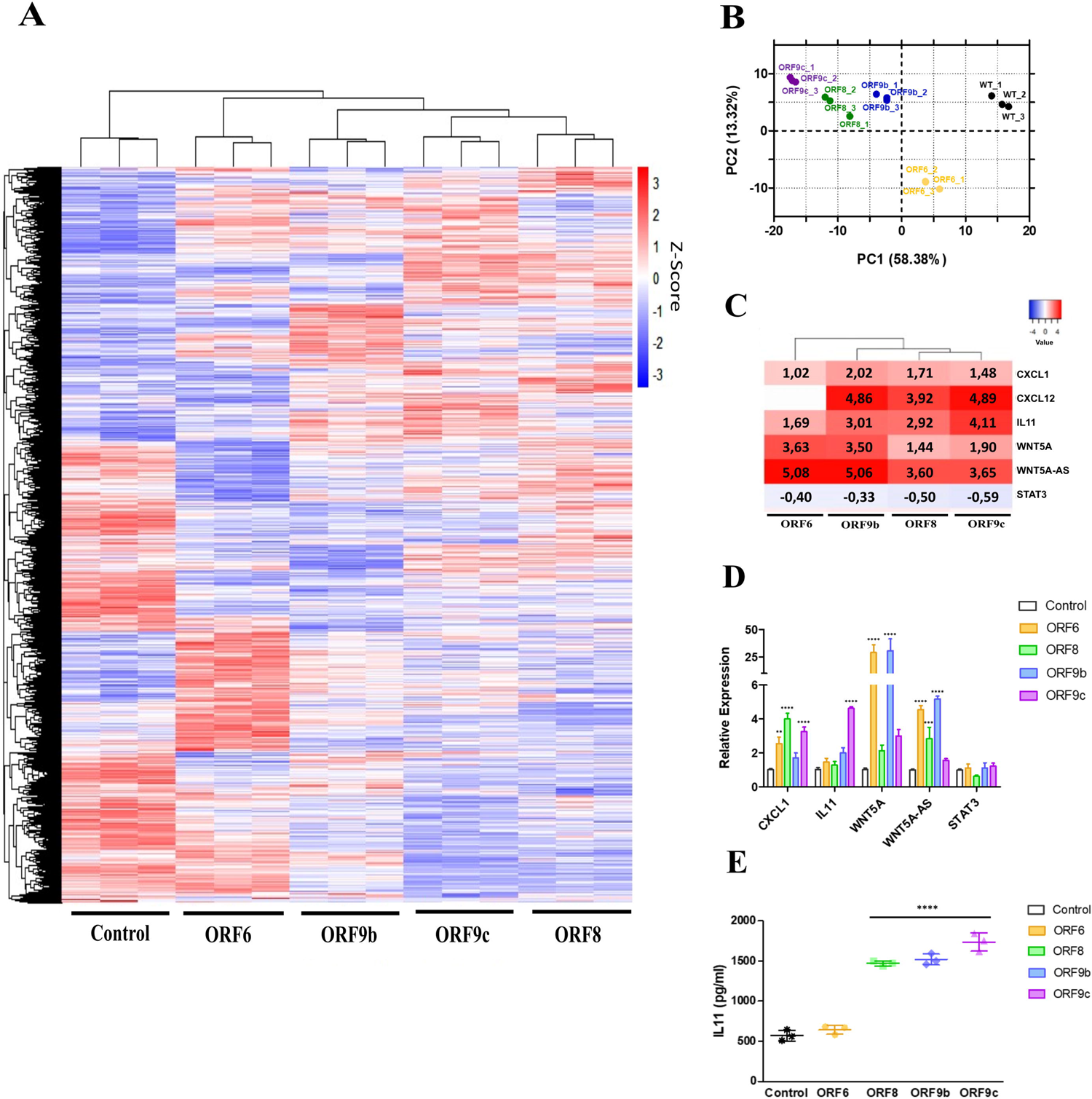
Differentially expressed genes (DEGs) in ORF-A549. **A)** Heatmap of RNA-Seq analysis of transduced cells expressing viral proteins. **B)** PCA graph of A549 control cells and A549 cells transduced with ORF6, ORF8, ORF9b or ORF9c. **C)** Log2 Fold Change heatmap of WNT5A and IL11 signaling pathways related genes. **D)** qRT-PCR gene expression levels calculated with 2-ΔΔCT method by normalizing to that of GADPH. **E)** ELISA of secreted IL11 by transduced cells after 24h. Error bars represent mean ± SD (n=3). Statistical significance is given as follows: *pc<c0.05, **pc<c0.01, ***pc<c0.001, **** pc<c0.0001 to A549 control cells.

### A549 transduced cells differently express genes involved in Pulmonary Fibrosis Idiopathic Signaling

Based on the above results, a functional pathway analysis with Ingenuity Pathway Analysis (IPA) software was performed. Both WNT5A and IL11 related canonical pathways were selected and two canonical pathways were found in common between the four transduced cell lines: Cardiac Hypertrophy Signaling and Pulmonary Fibrosis Idiopathic Signaling (Figure 3A). Genes involved in pulmonary fibrosis of each transduced cell line were obtained and further analysis showed a high gene expression pattern similarity between ORF-A549 cells (Figure 3B). Subsequent qRT-PCR experiments corroborated differential gene expression for collagen genes such as *COL1A1, COL4A1* or *COL11A1*, or other genes like *ADAMTS1, BCL2, IL1B, MMP16*, *SERPINE1, SNAI1* or *TFGB1* (Figure 3C). To assess whether altered expression of these genes could affect the profibrotic behavior of the cells, a functional assay was performed to test the ability of ORF-A549 cells to contract a collagen matrix (Figure 3D). After 24h, ORF6 and ORF9b expressing cells were able to significantly shrink the collagen matrix, while ORF8 and ORF9c transduced cells were able to do it only after 48h. Surprisingly, the contractile capacity of ORF9b-A549 cells was significantly higher than the others.

**Figure 3.**
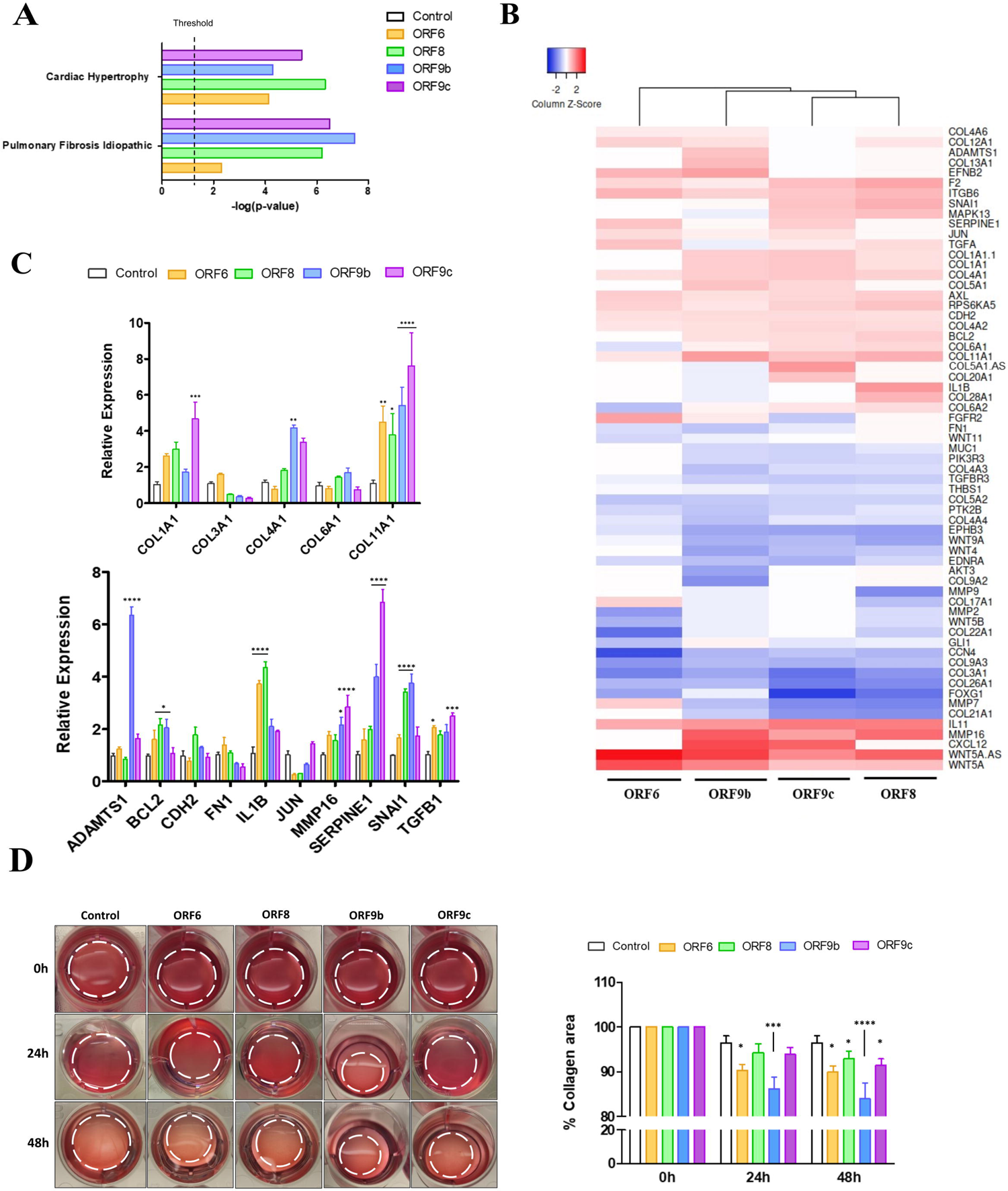
Pulmonary fibrosis idiopathic signaling pathway genes in ORF-A549. **A)** Both common WNT5A and IL11 related canonical pathways in transduced cells (IPA software analysis). **B)** Differentially expressed genes involved in pulmonary fibrosis of each transduced cell lines. **C)** qRT-PCR gene expression levels of various common genes calculated with 2-ΔΔ CT method by normalizing to that of GADPH. **D)** Representative cell contraction assay showing the ability of cells to shrink a collagen matrix in vitro. Dashed lines designate the gel edges. Bars indicate quantification of % collagen area contraction. Data are represented as mean ± SD (n=3). Statistical significance is given as follows: *p < 0.005, **p < 0.01, ****p < 0.0001 to A549 control cells.

### Inhibition of IL11 signaling pathway modulates the effect of ORF6, ORF8, ORF9b and ORF9c expression in A549 cells

Involvement of IL11 with fibrosis has been shown in previous studies ^31, 35, 37, 55^. To decipher IL-11 involvement, ORF-A549 cells were treated with an IL11 receptor inhibitor: Bazedoxifene (BAZ). ORF-A549 cells were treated with 5 μM BAZ during 24h and expression levels of various genes involved in fibrosis were determined (Figure 4, A-G). A decrease in *IL11* expression levels was observed in ORF8, ORF9b and ORF9c expressing cells, but not in ORF6-A549 cells (Figure 4A). These results were in agreement with those observed for IL11 secretion measured by ELISA (Figure 4H). As expected, we found changes in *WNT5A* after BAZ treatment (Figure 4B), but they were cell dependent. After IL11 signaling inhibition by BAZ, only ORF8-A549 cells showed a decrease in *WNT5A* expression. Interestingly, ORF8-A549 cells had the smallest increase in *WNT5A* when validated by RT-qPCR (Figure 2D). By contrast, ORF9b-A549 cells increased *WNT5A* expression after BAZ treatment, but no changes were observed in ORF6 or ORF9c expressing cells. Surprisingly, we did not observe any change in *TGFβ* expression in any ORF-A549 cells (Figure 4C). These results suggest that IL11 involvement in such profibrotic processes might be not mediated by TGFβ signaling. A decrease in *SERPINE1* expression after BAZ treatment was observed, particularly in ORF8, ORF9b and ORF9c expressing cells (Figure 4D). On the other hand, a significant increase in *IL1B, SNAI1* and *ADAMTS1* expression was observed in ORF9b-A549 cells after BAZ treatment (Figure 4, E-G). No changes in expression of these genes were shown in ORF6, ORF8 or ORF9c cell lines, suggesting a crosslink between IL11 and IL1B signaling pathways in ORF9b-A549 cells.

**Figure 4.**
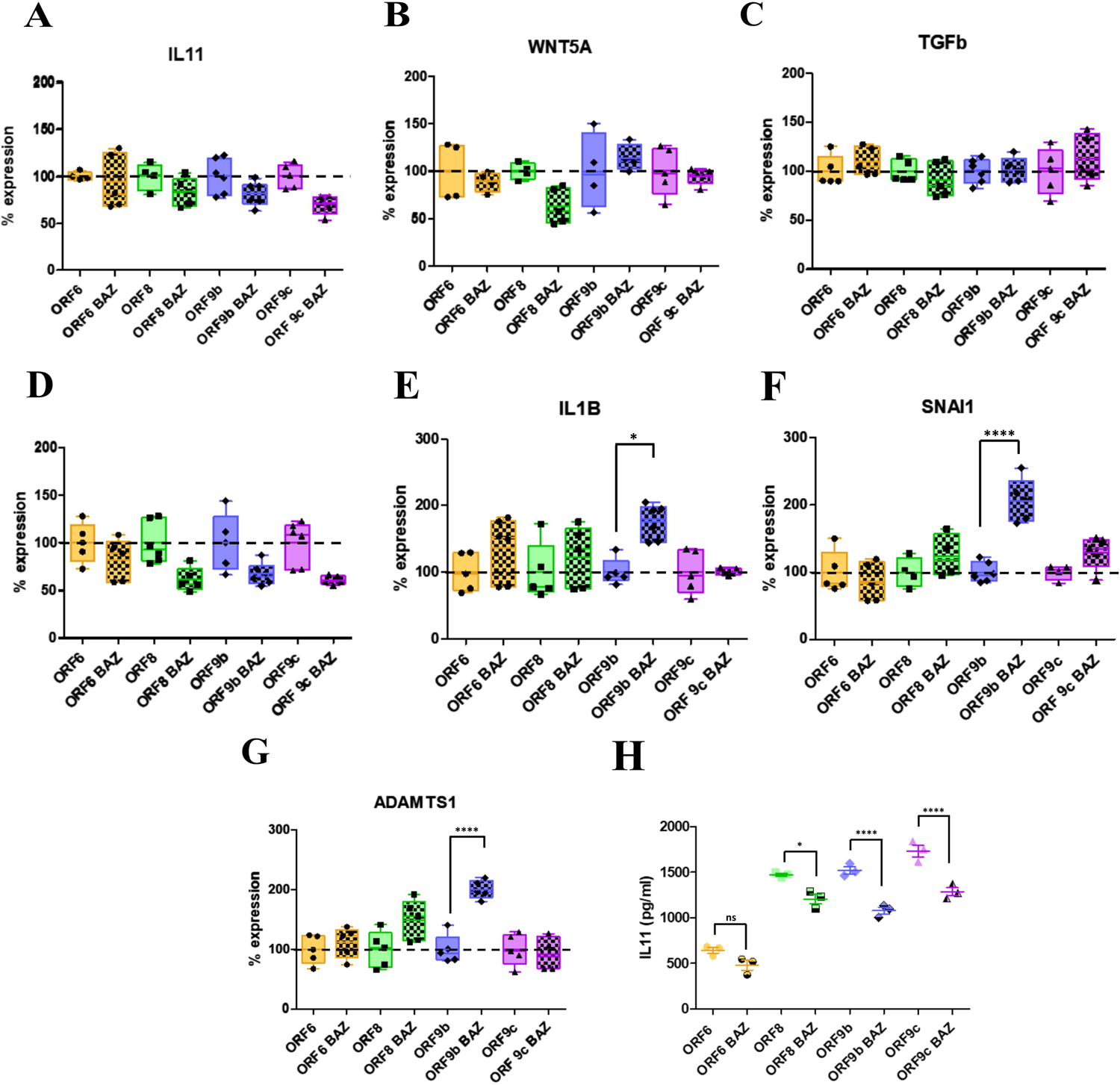
Alterations in gene expresión and IL11 release after 5µM Bazedoxifene (BAZ) 24h treatment. IL11 **(A),** WNT5A **(B),** TGFb **(C),** SERPINE1 **(D),** IL1B **(E),** SNAI1 **(F)** and ADAMTS1 **(G)** expression levels in BAZ treated cells compared to untreated cells. **H) ELISA** of IL11 secreted by transduced cells treated with BAZ compared to untreated cells. Data are represented as mean ± SD (n=3). Statistical significance is given as *p < 0.05 or ****p < 0.0001 to untreated cells.

IL11 increase after viral infections ^33, 34^ and a relationship between IL11 and WNT5A through STAT3 pathways signaling has been previously described ^40^. Therefore, STAT3 phosphorylation after IL11 signaling inhibition was analysed by western blot. A significant reduction in STAT3 phosphorylation was observed in cells expressing ORF8 and ORF9c (Figure 5A and 5B). It is reported that activation of the TGFβ signaling cascade causes phosphorylation and activation of the cytoplasmic effectors such as Smad2 ^59^. However, we did not observe changes in TGFβ expression nor Smad2 phosphorylation in any ORF-A549 cells (Figure 5A, 5C and 5D). These results were consistent with those observed by qRT-PCR (Figure 4C), where no changes in TGFβ expression were observed. Once more, these results suggest that IL11 involvement in this process may not be TGFβ dependent. Surprisingly, we did not observe significant changes in WNT5A expression after BAZ treatment (Figure 5, E-F). A significant decrease of WNT5A expression was found in ORF9c-A549 cells, but BAZ treatment did not alter such expression (Figure 5F). Regarding SERPINE1, a reduction in its expression by cells expressing ORF6 and ORF9c after BAZ treatment was observed, but it was only significant in ORF6-A549 (Figure 5E and 5G). Interestingly, a significant increase of phosphorylated c-jun in cells expressing ORF9b and ORF9c was found. In addition, BAZ treatment reduced phosphorylated c-jun in these cell lines (Figure 5H). However, ORF6 and ORF8 cell lines did not show changes in phosphorylated c-jun, and even BAZ treatment significantly augmented phosphorylated c-jun in cells expressing ORF6.

**Figure 5.**
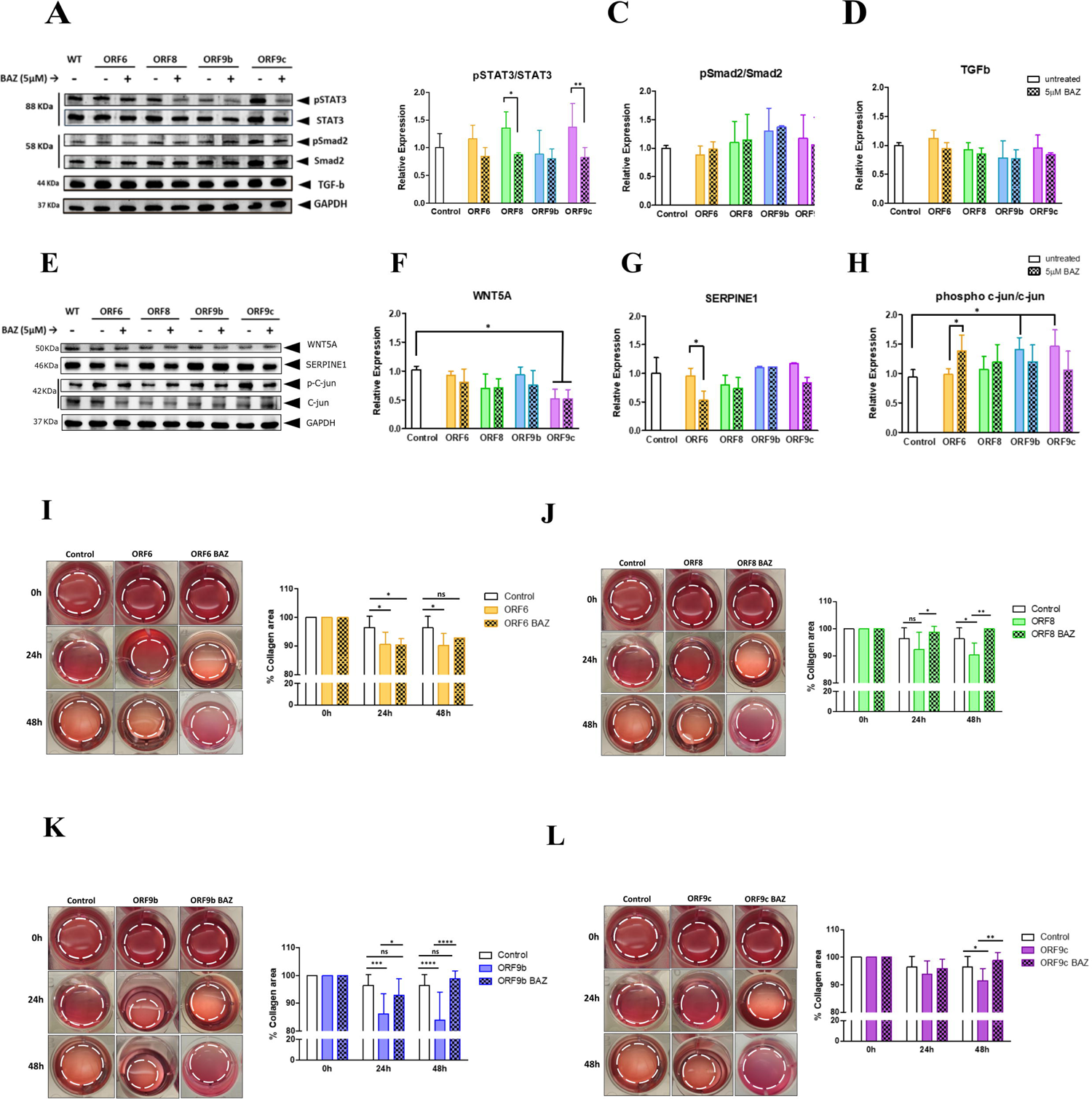
Effect of IL11 signalling inhibition by 5µM Bazedoxifene (BAZ) in protein expression and collagen gel contraction. **A)** Western blot of pSTAT3, STAT3, pSmad2, Smad2 and TGFb in cells treated 24h with BAZ compared to untreated cells. Ratio of phosphorylated/non-phosphorylated STAT3 **(B)** and Smad2 **(C). (D)** TGFb expression quantification. **E)** WNT5A, SERPINE1, phospho Ser73 c-jun and c-jun protein expression by Western Blot. WNT5A **(F),** SERPINE1 **(G)** and ratio of phosphorylated/non-phosphorylated c-jun **(H)** expression quantification. Statistical significance is given as *p < 0.05 or **p < 0.01. I-L: Representative cell contraction assay of A549 transduced cells treated with 5µM Bazedoxifene after 24h and 48h: ORF6 **(I),** ORF8 **(J),** ORF9b **(K)** and ORF9c **(L).** Statistical significance is given as follows: *p < 0.0005, **p < 0.01 ***p < 0.001 ****p < 0.0001. In all cases data are represented as mean ± SD (n=3).

Given the fact that expression of these accessory proteins modified their profibrotic capacity, IL11 involvement was analysed by BAZ treatment inhibiting IL11 signaling pathway in the collagen contraction assay (Figure 5, I-L). Interestingly, all ORF-A549 cells were able to revert the effect of expressing ORF6, ORF8, ORF9b or ORF9c accessory proteins. After 24h of treatment, we did not find changes in ORF6 and ORF9c cells compared to control cells (Figure 5I and 5L), but we did in cells expressing ORF8 and ORF9b (Figure 5J and 5K). By contrast, after 48h of BAZ treatment, all ORF-A549 cells recovered similar levels of collagen area when compared with untreated control cells. Therefore, these data indicate that IL11 signaling pathway is directly related to the profibrotic capacity described in ORF-A549 cells.

### Profibrotic response of lung epithelial cells to SARS-CoV-2 accessory proteins resemble responses to whole virus infection

In order to investigate the relevance of these profibrotic processes in SARS-CoV-2 virus infection, a bioinformatics comparative study was performed by integrating transcriptomic results from SARS-CoV-2 infected lung cell lines or COVID19 lung biopsies with those obtained in this study. The aim was to analyse common genes differentially expressed and their possible relationship with host fibrotic response when the whole virus was present. To this end, we grouped the sets of fibrosis-related genes in ORF-A549 cells obtained by IPA analysis, and a single common list of 63 fibrosis-related genes was generated (Figure 3B). Subsequently, our differential expression data list was compared with those obtained from infecting ACE2-transfected A549 cells and Calu3 cells with SARS-CoV-2 (NCBI-GEO, GSE147507) ^60^ (Figure 6A). Interestingly, we found 4 common genes between the three lung cell lines (*IL11, SNAI1, COL4A1* and *COL4A2*), as well as 4 common genes between our ORF-A549 cells and ACE2-transfected A549 cells (*COL11A1, COL21A1, COL5A2* and *COL6A1*), and 3 common genes between our ORF-A549 cells and Calu3 cells (*SERPINE1, THBS1* and *MUC1*). Gene expression disclosed two genes commonly upregulated (*IL11* and *SNAI1*) among lung cell lines, except in the case of ORF6-A549 cells, where *SNAI1* was not differentially expressed (Figure 6A).

**Figure 6.**
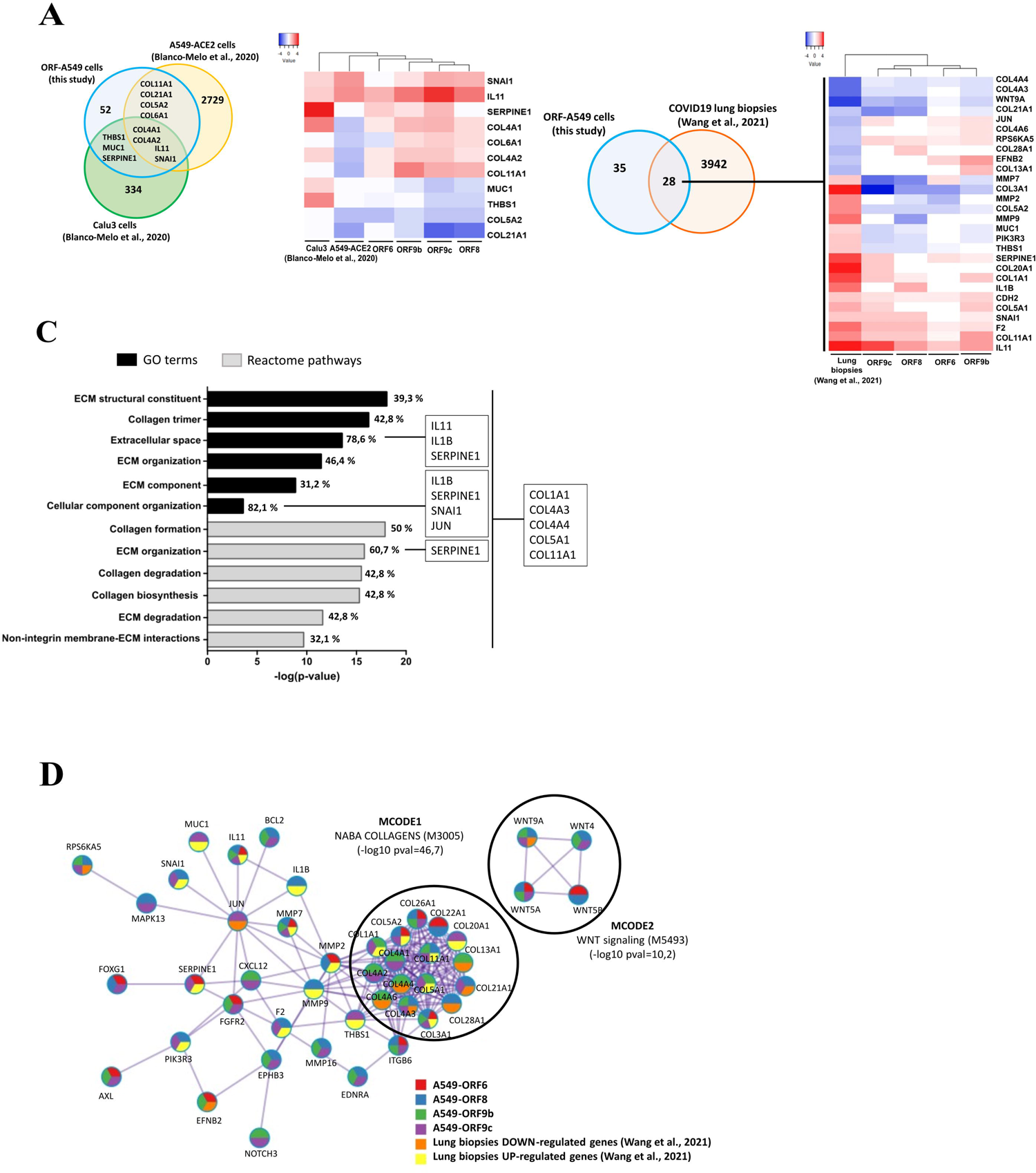
Comparison of gene expression responses with SARS-CoV-2 infected lung cell lines and COVID19 lung biopsies. **A)** Venn diagram of the intersection between differentially expressed fibrosis-related genes in ORF-A549 cells generated in this study with two SARS-CoV-2 infected lung cell lines from Blanco-Melo et al., 2020 (left) and heatmap showing differential gene expression pattern (right). **B)** Venn diagram of the intersection between differentially expressed fibrosis-related genes in ORF-A549 cells generated in this study with COVID19 lung biopsies from Wang et al., 2021 (left) and heatmap showing differential expression pattern (right). **C)** Enrichment study clustering common fibrosis-related genes between ORF-A549 cells generated in this study with COVID19 lung biopsies according to GO Terms and Reactome pathways. Most statistically significant pathways involved in fibrosis are represented. Percentage of genes implicated in each category is indicated in each bar. More representative genes in one or all pathways are stated. **D)** Most relevant MCODE components identified from the PPI network. Network nodes are displayed as pies. The color code represents a gene list (metascape.org).

**Figure 7.**
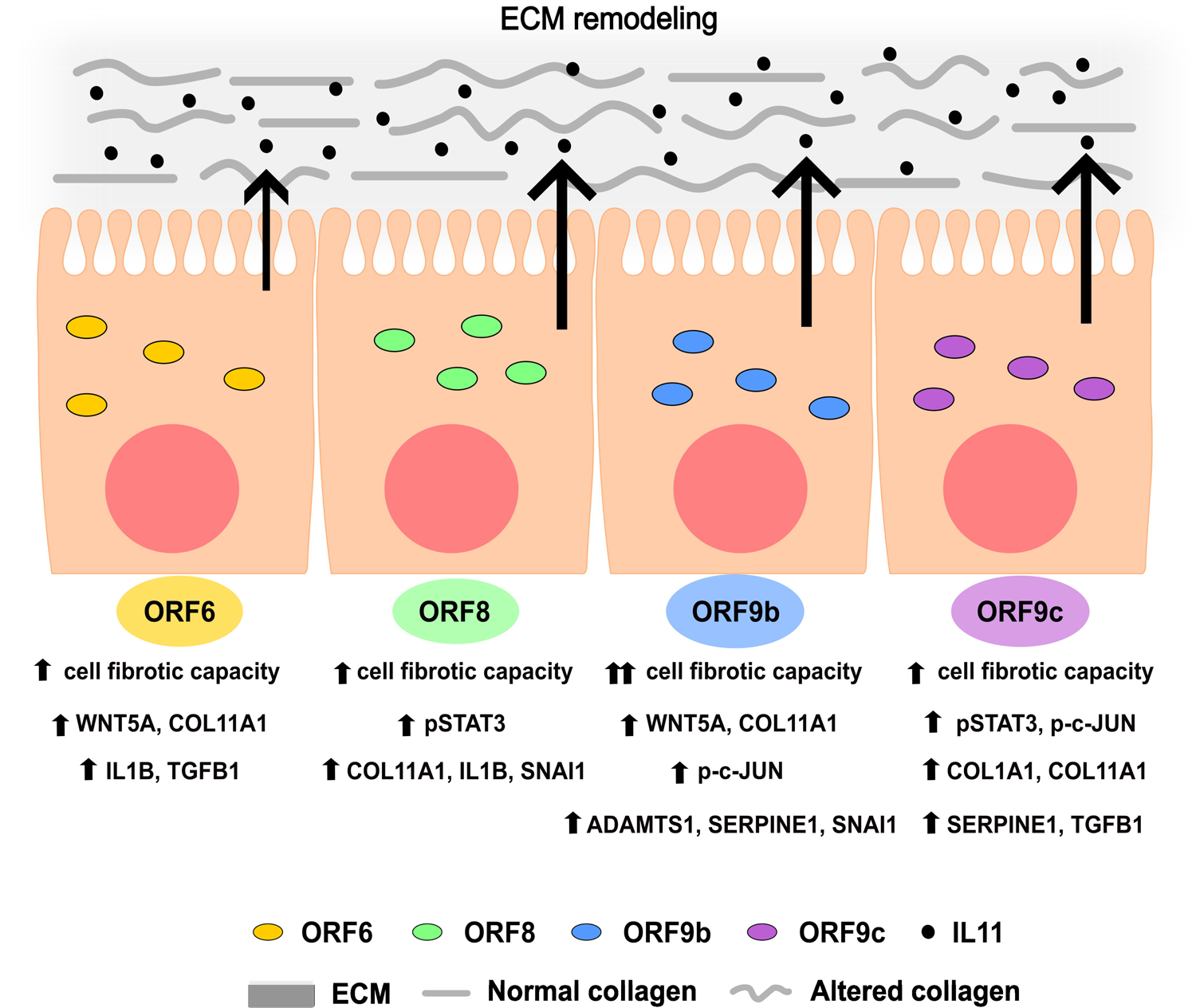
Graphical summary. Effects of SARS-CoV-2 accessory proteins ORF6, ORF8, ORF9b or ORF9c in A549 lung epithelial cells described in this study.

When we compared our list of fibrosis-related genes with transcriptomic data from post-mortem COVID19 lung biopsies (https://github.com/Jiam1ng/COVID-19_Lung_Atlas) ^61^, 28 common genes among two data lists were found (Figure 6B). Further analysis of gene expression revealed 4 genes commonly downregulated (*COL4A4, COL4A3, WNT9A* and *COL21A1*) among lung biopsies and ORF-A549 cells, except in the case of ORF6-A549, where *COL4A3* and *COL21A1* were not differentially expressed. At the same, 10 genes were found commonly upregulated, nevertheless, only four of them were upregulated by ORF6-A549 cells (*SERPINE1*, *CDH2, F2* and *IL11*). Once again, ORF6-A549 cells had the fewest genes in common with the other cell lines and lung biopsies. These data were in agreement with those previously shown, for example, in terms of heatmap clustering, PCA or IL11 secretion, which show the difference of ORF6-A549 with the rest of ORF-A549 cells (Figure 1A, 1B and 1E).

To further investigate differential perturbation of pathways regulated by ORF6, ORF8, ORF9b and ORF9c accessory proteins in SARS-CoV-2 infection, the 28 common genes list was used to perform an enrichment study with DAVID Functional Annotation Tool, where selected genes were clustered according to GO Terms and Reactome pathways (Figure 6C). We obtained the most statistically significant pathways involved in fibrosis and calculated the percentage of genes from the 28 common genes list. As expected, a clear predominance of terms and pathways related with ECM remodeling was observed. They were ECM structural constituent (GO:0030020), extracellular space (GO:0005615), ECM organization (GO:0030198 and R-HSA-1474244), ECM component (GO:0044420), cellular component organization (GO:0016043), ECM degradation (R-HSA-1474228) or non-integrin membrane-ECM interactions (R-HSA-3000171). Similarly, various terms or pathways implicated with collagen formation were disclosed, such as collagen trimer (GO:0005581), collagen formation (R-HSA-1474290), collagen degradation (R-HSA-1442490) and collagen biosynthesis (R-HSA-1650814) (Figure 6C). These results were in line with preliminary results obtained by proteomics analysis, in which three enzymes involved in the formation of collagen fibers were found altered (data not shown). All represented terms showed 5 common genes between our ORF-A549 cells and COVID19 lung biopsies (*COL1A1, COL4A3, COL4A4, COL5A1* and *COL11A1*). Apart from that, other commonly upregulated genes were found in certain terms, such as *IL11, IL1B, SERPINE1, SNAI1* and *JUN*. Interestingly, both cytokines, IL11 and IL1B were localized in extracellular space. Later, additional enrichment analysis and PPI network were obtained with MCODE network components (Metascape) (Figure 6D). Top two best p-value terms were retained: MCODE1, related to genes encoding collagen proteins (NABA_COLLAGENS, M3005, -log10 pval=46,7), and MCODE2, related to WNT signaling (M5493, -log10 pval=10,2). Although we did not observe any MCODE component clustering genes such as *IL11, IL1B, SERPINE1* or *SNAI1*, we did notice a relationship between these genes in the PPI network. Interestingly, *JUN* appeared as a connecting node of this cluster of genes.

## Discussion

SARS-CoV-2 virus, responsible for COVID19 disease, is associated with extensive lung alterations which can derive in pulmonary fibrosis ^5^. Indeed, recent bibliography has confirmed COVID19-fibrotic alterations ^1–4, 62^, which are even presented in long-COVID19 patients during the first year following the virus infection ^6^. In this study, A549 lung epithelial cells were individually transduced with accessory proteins ORF6, ORF8, ORF9b or ORF9c from SARS-CoV-2 (Wuhan-Hu-1 isolate), and subsequent transcriptomic with bioinformatic analysis disclosed that these accessory proteins can be involved in inflammatory and/or fibrotic responses in SARS-CoV-2 infection.

Noteworthy, virulent strains such as MERS-CoV, SARS-CoV and SARS-CoV-2 have a significant number of these accessory proteins, while more harmless coronaviruses have less ^63, 64^. This suggests that accessory proteins play a key role in pathogenesis not observed in less virulent coronavirus infections, although they have been less characterized than other proteins contained in the viral genome. Importantly, mutations in accessory proteins ORF6, ORF8 and ORF9b have been observed in currently circulating SARS-CoV-2 “variants of concern”, thus potentially contributing to increasing pathogenesis and transmissibility (https://covariants.org/variants).

At the beginning of the pandemic, high levels of IL6 in COVID19 patient serum were described to correlate with severe disease ^50, 51^. Surprisingly, we found a high overexpression of IL11 in epithelial transduced cells (Figures 2C and 2D), while no changes in IL6 expression or release were observed (data not shown). Indeed, IL11 release was also augmented in three of the four transduced cell lines (Figure 2E). These results agree with those reported in several studies, where IL11 was defined as an “epithelial interleukin”, while IL6 biology was related mostly to immune functions^25, 29, 31^. In addition, several studies have related high levels of IL11 with fibrosis, chronic inflammation and matrix extracellular remodeling ^31, 35–39^. However, whether this elevation is pathogenic or a natural host response to restore homeostasis remains unanswered for many diseases ^36^.

We also found several fibrosis related genes differentially expressed. Among them, *WNT5A* was particularly upregulated in ORF6 and ORF9b expressing cells. Likewise, we found an increase in *WNT5A-AS* (Figures 2C and 2D). WNT5A is a member of WNT family proteins which plays critical roles in a myriad of processes in both health and disease ^41^, and it is known its relationship with IL11 through STAT3 pathways signalling ^40^. Besides, chemokines CXCL1 and CXCL12 have been found to be upregulated by WNT5A in various studies ^41, 52^. These results are consistent with those we have observed in transcriptomic analysis, where both chemokines were upregulated together with *WNT5A* (Figures 2C and 2D), although they did not exactly correlate with ORF6 and ORF9b expressing cells. WNT5A-AS has been reported as a long noncoding RNA (lncRNA) located on the antisense strand of chromosome 16 p16, and which overlaps with introns of *WNT5A* on the sense strand ^65^. Indeed, Lu et al. and Salmena et al. provided evidence of a positive correlation between the upregulation of lncRNA WNT5A-AS with that of its antisense gene, *WNT5A*, suggesting that lnRNA WNT5A-AS acts as a competing endogenous RNA to regulate the expression of *WNT5A* ^65, 66^. In this study, we found high levels of *WNT5A* gene expression in ORF6 and ORF9b expressing cells, but we did not observe this increase in protein expression (Figure 5E and 5F). Thus, it is plausible to think that *WNT5A-AS* was responsible for regulating posttranscriptional expression of *WNT5A*.

TGFβ represents the most prominent profibrotic cytokine by upregulating production of ECM components and multiple signaling molecules ^47^. There is, indeed, a clear evidence of the relationship between IL11, WNT, TGFβ and fibrosis ^42–46^. In this study, *TGFβ* was among the genes related to fibrosis, and we observed an increase in *TGFβ* gene expression but our assays did not find any upregulation in its protein expression (Figures 5A and 5D). When canonical TGFβ signaling pathway through Smad2 phosphorylation was analysed, no significant changes were also observed (Figures 5A and 5C), indicating that TGFβ involvement may be regulated either by non-canonical TGFβ signaling pathway or through TGFβ-independent manner.

Within the list of genes involved in fibrosis, we also found genes such as *ADAMTS1, BCL2, CDH2, FN1, IL1B, JUN, MMP16, SERPINE1, SNAI1* and various collagen genes (Figures 3B and 3C). Expression of these genes differed depending on the expressed accessory protein. In organs, such as the lungs, resident cells actively and continuously remodel the extracellular matrix (ECM), forming a dynamic network balanced by cell–ECM bidirectional interactions ^67^. In order to check how A549 transduced cells responded to and actively remodeled the ECM, we performed a collagen gel contraction assay. A strong ability in ORF9b-A549 cells and moderate ability in ORF6-A549 cells to contract the collagen gel in the first 24 hours was observed, which was also followed by ORF8 and ORF9c transduced cells after 48 hours. These data indicate that expression of these accessory proteins, and particularly ORF9b, trigger a profibrotic process. In addition, preliminary proteomics analysis revealed three altered enzymes involved in collagen fibers formation: PLOD1, PLOD2 and COLGALT1 (data not shown). It is known that PLOD1 and PLOD2 catalyze the lysyl hydroxylation to hydroxylysine, which is critical for the formation of covalent cross-links and collagen glycosylation ^53^. COLGALT1 acts on collagen glycosylation and facilitates the formation of collagen triple helix ^54^, and an increase of *WNT5A* gene expression has been recently correlated with COLGALT1 downregulation ^68^. These three enzymes tend to be downregulated in ORF-A549 cells, except in ORF6-A459 cells, where PLOD2 was significantly increased. Furthermore, PLOD1 was significantly decreased in ORF8-A549 cells, while PLOD2 and COLGALT1 were significantly decreased in ORF9b-A549 cells, meaning a possible involvement of these enzymes in the increased collagen-contraction ability of these cells (data not shown).

Our hypothesis was that IL11 might be behind the above mentioned profibrotic alterations in ORF-A549 cells, so we used an IL11 receptor inhibitor to block IL11 signaling pathway. Several studies have identified BAZ as a novel small-molecule inhibitor of GP130 ^56^, and support its therapeutic action targeting IL-11/GP130 signaling for cancer therapy ^57, 58^. BAZ binds to GP130 heterodimer and inhibits IL6 family members-induced STAT3 phosphorylation ^56^, blocking interleukins signaling pathways without affecting their release, as we observed (Figure 4H). In this study, BAZ treatment of ORF-A549 cells reverted their high collagen-contraction ability (Figure 5, I-L). STAT3 phosphorylation was also reduced in ORF8 and ORF9c expressing cells (Figure 5A and 5B). Noteworthy, ORF8 and ORF9c expressing cells showed the lowest levels of *WNT5A* gene expression. These data suggest a possible IL11 signaling pathway regulation by *WNT5A*. We also observed a decrease in IL11 release by these cells, but we did not notice a significant downregulation of any genes altered by accessory proteins expression (Figure 4). Interestingly, an increase in *IL1B, SNAI1* and *ADAMTS1* in ORF9b expressing cells after BAZ treatment was found, suggesting a possible crosslink between IL11 and IL1B signaling pathways. Palmqvist et al. provided evidence of enhanced IL11 expression by IL1B by a mechanism involving MAPK in gingival fibroblasts ^69^. Nevertheless, further investigations must be performed to test this hypothesis and its relationship with high levels of *SNAI1* and *ADAMTS1* when blocking IL11 signaling pathway.

Finally, data comparison with lung cell lines infected with SARS-CoV-2 and lung biopsies from patients with COVID19 showed evidence of altered gene expression that matched with results obtained in this study. Firstly, we found common differentially expressed genes with SARS-CoV-2 infected lung cell lines ^60^. Among these genes, *IL11* was commonly upregulated, as well as *SNAI1*. On the other hand, 28 common genes related with fibrosis were found between our transduced cells lines and COVID19 lung biopsies ^61^. We also found *IL11* between this cluster of genes. A subsequent enrichment analysis showed that this set of genes was mostly involved in ECM organization and collagen formation (Figure 6C). Interestingly, both IL11 and IL1B were located in extracellular space. These data were consistent with the fact that we did not find these interleukins in proteomics study or by western blot assay (data not shown). Five collagen genes were common in all signaling pathways (*COL1A1*, *COL4A3, COL4A4, COL5A1* and *COL11A1*) and they were also clustered together by MCODE algorithm using Metascape tool (Figure 6D). Remarkably, *JUN* was listed in the cellular component organization GO term (Figure 6C), and it was also found in PPI network as a gene connecting node (Figure 6D). That pointed to a possible c-jun role connecting profibrotic cell responses. Ser73 c-jun phosphorylation was confirmed by western blot in ORF9b and ORF9c expressing cells (Figure 5E and 5H). Indeed, this c-jun activation decreased after BAZ treatment. Noteworthy, phosphorylation of c-jun increased in ORF6-A549 cells after blocking IL11 signaling pathway, indicating that c-jun activation in ORF9b-A549 and ORF9c-A549 cells was mediated by IL11 expression. C-jun Ser73 is phosphorylated by MAPK8 ^70^, and JNK-interacting proteins (JIP) are a scaffold proteins group that selectively mediates JNK signaling by aggregating specific components of the MAPK cascade. Among JIP proteins, SPAG9 or JIP4 (also known as MAPK8IP4) is involved in MAPK signaling pathway to regulate cellular activities^71^. Del Sarto et al. recently have identified an increase of phosphorylaton at Ser730 in JIP4 after Influenza A virus infection ^72^. Similarly, other work has recently described that this phosphorylation promotes cell death via c-jun kinase signaling pathway ^73^. Thus, there is a correlation between c-jun phosphorylation and SPAG9 (JIP4) phosphorylation after Influenza A virus infection. Preliminary phosphoproteomics analysis in this study revealed the presence of phosphorylated form of SPAG9 in the position Ser730 (data not shown). However, further mass spectrometry analysis will be required to define the phosphorylation patterns and abundance changes of phosphorylated SPAG9 in ORF-A549 cells.

On the other hand, ORF6-A549 cells showed the lowest levels of IL11 release, differentially of the rest of ORF-A549 set of cells. Data from ORF6-A549 appeared separated from the other ORF-A549 cells in transcriptomic clusters (Figure 2B) and showed less genes in common when compared with SARS-CoV-2 infected lung cell lines and COVID19 lung biopsies (Figure 6A and 6B). Indeed, only ORF6-A549 cells showed an increase in PLOD2 enzyme, and a decrease in SERPINE protein expression after BAZ treatment. All these data indicate a different mechanism of action by ORF6 that require further investigations.

Taken together, our findings indicated that SARS-CoV-2 accessory proteins ORF6, ORF8, ORF9b and ORF9c have the ability to trigger profibrotic cell responses in A549 human lung epithelial cells. Interestingly, increased IL11 led to ECM remodeling. SARS-CoV-2 infected lung cell lines and COVID19 lung biopsies from patients show a similar response to SARS-Cov-2 infection, so these profibrotic responses may underlie COVID19-fibrotic alterations. Thus, these accessory proteins could be used as a target for new therapies for COVID19 disease against pulmonary fibrosis.

## Supporting information

Table S1_IL11-related IPA Canonical Pathways in ORF-A549 cells_related to Fig 3

Table S2_WNT5A-related IPA Canonical Pathways in ORF-A549 cells_related to Fig 3

Table S3_ORF-A549 cells and COVID19 lung biopsies comparison_DAVID Enrichment Study_related to Fig 6C

Table S4_Metascape analysis between ORF-A549 cells and COVID19 lung biopsies, related to Figure 6D

Table S5_List of Primers

## Acknowledgements

The authors wish to acknowledge Bioinformatics & Biostatistics Service at CIB and Dr Aurora Gómez-Durán (CIB).

## Author contributions

Conceptualization: M.M. and J.J.G.; Methodology: B.D.L.A., A.dL.R, L.M.G., T.G.G., R.F.R., J.M.S.C., F.M.S., F.C., N.R., F.P., S.Z.L. and A.J.M.; Investigation: B.D.L.A., A.dL.R, L.M.G., T.G.G., R.F.R., J.M.S.C., F.M.S., F.C., N.R., F.P., S.Z.L. and A.J.M.; Writing Original Draft: B.D.L.A.; Writing Review & Editing: M.M. and J.J.G.; Visualization: B.D.L.A.; Supervision: M.M. and J.J.G.; Project Administration: M.M. and J.J.G.; Funding Acquisition: M.M. and J.J.G.

## STARS Methods

### RESOURCE AVAILABILITY

#### Lead contact

Further information and requests for resources and reagents should be directed to and will be fulfilled by the lead contact, Dr. María Montoya González.

#### Materials availability

This study did not generate new unique reagents.

All unique material generated in this study are listed in the key resources table.

#### Data and code availability

All sequencing data sets are available in the NCBI BioProject database under accession number PRJNA946640 for A549 transduced cells and PRJNA841835 for A549 control cells.

## EXPERIMENTAL MODEL AND STUDY PARTICIPANT DETAILS

A549 pulmonary epithelial cells (ATCC CRM-CCL-185) were cultured in Dulbecco’s Modified Eagle Medium (DMEM) (Gibco) supplemented with 10% (v/v) heat-inactivated fetal bovine serum (FBS) (Gibco) and 1% Penicillin-Streptomycin (100U/ml) (Gibco). A549-transduced cells expressing SARS-CoV-2 ORF6, ORF8, ORF9b or ORF9c were additionally supplemented with 2 μ were cultured at 37°C in a 5% CO2, 90% humidity atmosphere.

### METHOD DETAILS

#### Lentivirus production and transduction

ORF6, ORF8, ORF9b or ORF9c coding sequences (codon-optimized for mammalian expression) were cloned into pLVX-EF1α-IRES-Puro Cloning and Expression Lentivector (Clontech, Takara) to generate pseudotyped lentiviral particles encoding the ORF6, ORF8, ORF9b or ORF9c accessory proteins of SARS-CoV-2 (Wuhan-Hu-1 isolate) at the CNIC (Centro Nacional de Investigaciones Cardiovasculares) Viral Vector Unit (ViVU), essentially as previously described ^74^. ORF6, ORF8, ORF9b or ORF9c accessory proteins were C-terminally 2xStrep-tagged to check viral protein expression. A549 pulmonary epithelial cells were transduced by incubating them with lentivirus at a MOI of 10 for 24 h followed by 2 µg/ml puromycin treatment to start the selection of successfully transduced cells. GFP expressing cells were generating by transducing them with pLVX-AcGFP1-N1 lentiviral particle (Clontech, Takara).

#### Immunofluorescence microscopy

Cells were seeded on 24-well plates containing glass coverslips coated with poly-lysine solution (100.000 cells per well). Cells were fixed with 4% PFA in PBS for 15 min, washed twice in PBS, and then permeabilized for 10 min with 0.1% Triton X-100 in PBS. Primary antibodies incubation was carried out for 1h in PBS containing 3% BSA and 0.1% Triton X-100 at 1:100 dilution. Coverslips were washed three times with PBS before secondary anti-mouse antibodies incubation (1:1000 dilution). The antibodies used for immunofluorescence are shown in the key resources table. Phalloidin was used as a cytoplasmic marker at 1:200, and DAPI (4’6-diamidino-2-phenylindole) (Molecular Probes) was used as a nuclear marker. Coverslips were mounted in Mowiol 4-88 (Sigma-Aldrich). Images were acquired with a confocal laser microscope Leica TCS SP8 STED 3X.

#### RNA isolation and sequencing

Cells were seeded (3 x 10^5^) in 6-well plates and lysed using RLT buffer for RNA isolation (RNeasy mini kit, Qiagen). Each sample was performed in triplicate. RNA was isolated following manufactureŕs protocol, quantified by nanodrop 1000 (Thermo Scientific) and quality controlled by Bioanalyzer (Agilent). All samples sent for sequencing had a RIN (RNA integrity number) over 9.90. cDNA libraries and sequencing were performed by Novogene Europe, using 400 ng of RNA per sample for library preparation. Samples were sequenced in an Illumina platform using a PE150 strategy.

#### Gene sets and differential gene expression analysis

Sequencing raw data was quality controlled (error rate, GC content distribution) and filtered, removing bad quality and N-containing sequences and adaptors. Clean data were mapped (HISAT2) to reference genome GRCh38.p13, and gene expression was quantified using FPKM (Fragments Per Kilobase of transcript sequence per Millions of base pairs sequenced). Differential expression analysis was performed using DESeq2 R package ^75^.

Raw counts were transformed with the vst function in the DESeq2 package ^76^ of the R software version 3.6.3 ^77^, and subsequent PCA was performed with the prcomp function. The 500 genes with the highest variance among samples were considered. Finally, the PCA graph was made with GraphPad Prism 5 (GraphPad software, San Diego, CA, USA).

#### Real time qPCR

RNA samples (500 ng) were reverse transcribed using qScript™ cDNA synthesis kit (Quanta Biosciences Inc.), following manufacturer’s instructions. Primers sequences are available in Supplementary Table 5 (Table S5). The final 15 µL PCR reaction included 2 μL of 1:5 diluted cDNA as template, 3 µL of 5x PyroTaq EvaGreen qPCR Mix Plus with ROX (Cultek Molecular Bioline, Madrid, Spain), and transcript-specific forward μM final concentration. Real time PCR was carried out in a QuantStudio 12K Flex system (Applied Biosystems) under the following conditions: 15 min at 95 °C followed by 40 cycles of 30 s at 94 °C, 30 s at 57 °C and 45 s at 72 °C. Melting curve analyses were performed at the end, in order to ensure specificity of each PCR product. Relative expression results were calculated using GenEx6 Pro software (MultiD-Göteborg, Sweden), based on the Cq values obtained.

#### Western blot

Transduced cells were harvested and lysed in ice-cold Pierce IP Lysis Buffer (#87787, Thermo Scientific) at 4° C. Cell lysates were mixed with 5× SDS-PAGE Sample Loading Buffer (MB11701, Nzytech) and heated at 95° C for 5 min. Protein samples were resolved by SDS polyacrylamide gel electrophoresis and transferred onto a PVDF membrane using Mini Trans-Blot System (1703935, Bio-Rad), followed by blocking for 1 h with 5% BSA in Tris-buffered saline-Tween20 buffer and probing with corresponding primary and secondary antibodies. The proteins were visualized by chemoluminiscence using ChemiDoc Imaging Systems (Bio-Rad). Relative protein expression was calculated by sequentially normalizing against the loading control (GAPDH).

#### Bazedoxifene Treatment and ELISA

Cells were seeded (3 x 10^5^) in 6-well plates and treated with 5 µM of Bazedoxifene acetate (PZ0018, Sigma) for 24h. To perform ELISA experiments, IL-11 levels in supernatants collected after 24 h treatment from different cell lines were detected with the Human DuoSet ELISA Kits (DY218) according to the manufacturer’s instructions.

#### Cell Contraction Assay

CytoSelect™ 24-well Cell Contraction Assay Kit (Cell Biolabs) was used according to the manufacturer’s instructions. Briefly, collagen gel lattice was prepared by mixing 4.5 x 10^6^ cells/mL with a collagen gel solution and added to each well of the 24-well cell contraction plate. After collagen polymerization, fresh media was added and wells were monitored for contraction over two days at 37°C and 5% CO2. The change in matrix diameter size (in millimeters) was determined with a ruler each 24h.

#### Bioinformatics Analysis. Database comparison. Pathway Enrichment Analysis, Network and PPI Module Reconstruction

Functional pathway analysis of transduced cells was performed with Ingenuity Pathway Analysis (IPA) software. Adjusted p-value less than 0.05 was considered as the cut-off criterion for pathway enrichment analysis. To compare our results in A549 lentivirus-transduced expressing individual viral accessory proteins ORF6, ORF8, ORF9b or ORF9c with whole virus-infected cell lines A549-ACE2 or Calu3, transcriptomic data from ^60^ were used. Subsequently, transcriptomic data from ^61^ were applied for patient samples comparison. Bioinformatics analysis were carried out making Venn diagrams with Venny 2.1 ^78^, and heatmaps with Heatmapper program. A further enrichment study was performed with DAVID Functional Annotation Tool where selected genes were clustered according to GO Terms and Reactome Gene Sets. Additionally, another pathway enrichment analysis and the gene network reconstruction were carried out using the online Metascape Tool (http://metascape.org) ^79^ with the default parameters set. Enrichment analyses were carried out selecting the genomics sources: KEGG Pathway, GO Biological Processes, Reactome Gene Sets, Canonical Pathways, and CORUM. Terms with p < 0.01, minimum count 3, and enrichment factor >1.5 were collected and grouped into clusters based on their membership similarities. P-values were calculated based on accumulative hypergeometric distribution, and q-values were calculated using the Benjamini-Hochberg procedure to account for multiple testing. To further capture the relationship among terms, a subset of enriched terms was selected and rendered as a network plot, where terms with similarity >0.3 are connected by edges. Based on Protein-Protein Interaction (PPI) enrichment analysis, we run a module network reconstruction based on the selected genomics databases. The resulting network was constructed containing the subset of proteins that form physical interactions with at least one other list member. Subsequently, by means of Molecular Complex Detection (MCODE) algorithm, we first identified connected network components, then a pathway and process enrichment analysis were applied to each MCODE component independently and the three best-scoring (by p-value) terms were retained as the functional description of the resulting modules.

## QUANTIFICATION AND STATISTICAL ANALYSIS

Statistical analyses were performed using GraphPad PRISM 5. P-values were determined using two-way ANOVA and Bonferroni test correction was applied. Unless otherwise stated, data are shown as the mean of at least three biological replicates. Significant differences are indicated as: *, p <0.05; **, <0.01; ***, p<0.001, ****, p<0.0001.

---

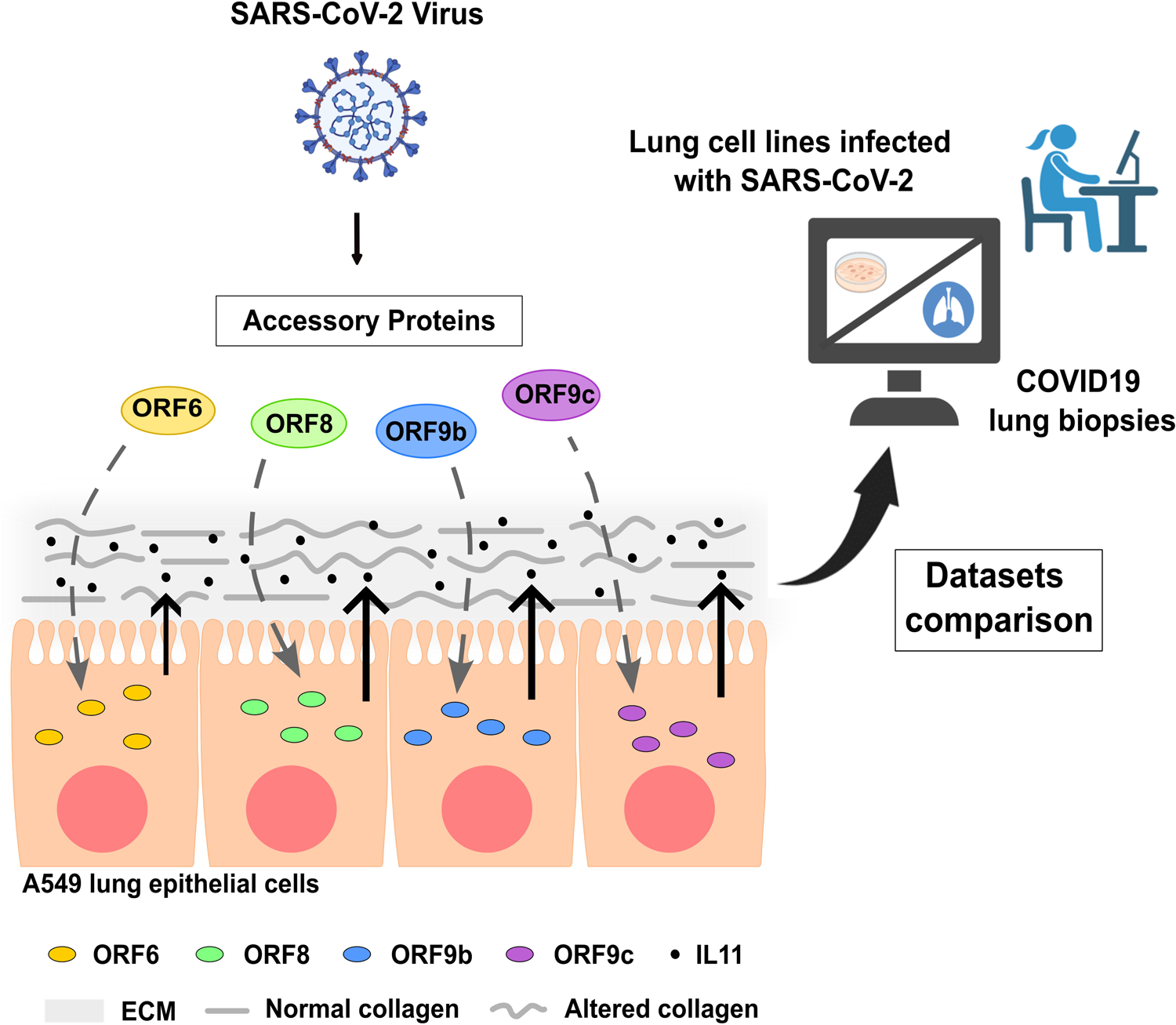

## References

1. Dinnon KH, Leist SR, Okuda K, et al. SARS-CoV-2 infection produces chronic pulmonary epithelial and immune cell dysfunction with fibrosis in mice. Sci Transl Med. 2022;14(664):1–33. doi:10.1126/scitranslmed.abo5070

2. John AE, Joseph C, Jenkins G, Tatler AL. COVID-19 and pulmonary fibrosis: A potential role for lung epithelial cells and fibroblasts. Immunol Rev. 2021;302(1):228–240. doi:10.1111/imr.12977

3. Bergantini L, Mainardi A, d’Alessandro M, et al. Common Molecular Pathways Between Post-COVID19 Syndrome and Lung Fibrosis: A Scoping Review. Front Pharmacol. 2022;13(March):1-13. doi:10.3389/fphar.2022.748931

4. Mohammadi A, Balan I, Yadav S, et al. Post-COVID-19 Pulmonary Fibrosis. Cureus. 2022;14(3):4–10. doi:10.7759/cureus.22770

5. Giacomelli C, Piccarducci R, Marchetti L, Romei C. Pulmonary fibrosis from molecular mechanisms to therapeutic interventions: lessons from post-COVID-19 patients. 2021;(January).

6. Fabbri L, Moss S, Khan FA, et al. Parenchymal lung abnormalities following hospitalisation for COVID-19 and viral pneumonitis: A systematic review and meta-analysis. Thorax. 2022;78(2):191–201. doi:10.1136/thoraxjnl-2021-218275

7. Ellis P, Somogyvári F, Virok DP, Noseda M, Mclean GR. Decoding Covid-19 with the SARS-CoV-2 Genome. Published online 2021:1–12.

8. Yoshimoto FK. The Proteins of Severe Acute Respiratory Syndrome Coronavirus-2 (SARS CoV-2 or n-COV19), the Cause of COVID-19. Protein J. 2020;39(3):198–216. doi:10.1007/s10930-020-09901-4

9. Redondo N, Zaldívar-López S, Garrido JJ, Montoya M. SARS-CoV-2 Accessory Proteins in Viral Pathogenesis: Knowns and Unknowns. Front Immunol. 2021;12(July):1–8. doi:10.3389/fimmu.2021.708264

10. Jungreis I, Nelson CW, Ardern Z, et al. Conflicting and ambiguous names of overlapping ORFs in the SARS-CoV-2 genome: A homology-based resolution. Virology. 2021;558(November 2020):145–151. doi:10.1016/j.virol.2021.02.013

11. Finkel Y, Gluck A, Nachshon A, et al. SARS-CoV-2 uses a multipronged strategy to impede host protein synthesis. Nature. 2021;594(7862):240–245. doi:10.1038/s41586-021-03610-3

12. Zhang Y, Chen Y, Li Y, et al. The ORF8 protein of SARS-CoV-2 mediates immune evasion through down-regulating MHC-. Proc Natl Acad Sci U S A. 2021;118(23):1–12. doi:10.1073/pnas.2024202118

13. Jiang H wei, Zhang H nan, Meng Q feng, et al. SARS-CoV-2 Orf9b suppresses type I interferon responses by targeting TOM70. Cell Mol Immunol. 2020;17(9):998–1000. doi:10.1038/s41423-020-0514-8

14. Xia H, Cao Z, Xie X, et al. Evasion of Type I Interferon by SARS-CoV-2. Cell Rep. 2020;33(1):108234. doi:10.1016/j.celrep.2020.108234

15. Miorin L, Kehrer T, Sanchez-Aparicio MT, et al. SARS-CoV-2 Orf6 hijacks Nup98 to block STAT nuclear import and antagonize interferon signaling. Proc Natl Acad Sci U S A. 2020;117(45):28344–28354. doi:10.1073/pnas.2016650117

16. Lee JG, Huang W, Lee H, van de Leemput J, Kane MA, Han Z. Characterization of SARS-CoV-2 proteins reveals Orf6 pathogenicity, subcellular localization, host interactions and attenuation by Selinexor. Cell Biosci. 2021;11(1):1–12. doi:10.1186/s13578-021-00568-7

17. Savellini GG, Anichini G, Gandolfo C, Cusi MG. Nucleopore Traffic Is Hindered by SARS-CoV-2 ORF6 Protein to Efficiently Suppress IFN-β and IL-6 Secretion. Viruses. 2022;14(6). doi:10.3390/v14061273

18. Wang X LJ et al. Accurate Diagnosis of COVID-19 by a Novel ImmunogenicnSecreted SARS-CoV-2 ORF8 Protein. 2020;11(5):1-13.

19. Beaudoin-Bussières G, Arduini A, Bourassa C, et al. SARS-CoV-2 Accessory Protein ORF8 Decreases Antibody-Dependent Cellular Cytotoxicity. Viruses. 2022;14(6):1–12. doi:10.3390/v14061237

20. Han L, Zhuang MW, Deng J, et al. SARS-CoV-2 ORF9b antagonizes type I and III interferons by targeting multiple components of the RIG-I/MDA-5–MAVS, TLR3– TRIF, and cGAS–STING signaling pathways. J Med Virol. 2021;93(9):5376–5389. doi:10.1002/jmv.27050

21. Faizan MI. NSP4 and ORF9b of SARS-CoV-2 Induce Pro-Inflammatory Mitochondrial DNA Release in Inner Membrane-Derived Vesicles. Cells. 2022;11(19). doi:10.3390/cells11192969

22. Dominguez Andres A, Feng Y, Campos AR, et al. SARS-CoV-2 ORF9c Is a Membrane-Associated Protein that Suppresses Antiviral Responses in Cells. bioRxiv Prepr Serv Biol. Published online 2020. doi:10.1101/2020.08.18.256776

23. Gordon DE et al. A SARS-CoV-2 Protein Interaction Map Reveals Targets for Drug-Repurposing. Gordon et al (Nevan). Nature Ap2020 FINAL. Nature. 2020;583:459-468. https://www.nature.com/articles/s41586-020-2286-9

24. Montazersaheb S, Hosseiniyan Khatibi SM, Hejazi MS, et al. COVID-19 infection: an overview on cytokine storm and related interventions. Virol J. 2022;19(1):1–15. doi:10.1186/s12985-022-01814-1

25. Cook SA, Schafer S. Hiding in Plain Sight: Interleukin-11 Emerges as a Master Regulator of Fibrosis, Tissue Integrity, and Stromal Inflammation. Annu Rev Med. 2020;71:263–276. doi:10.1146/annurev-med-041818-011649

26. Metcalfe RD, Putoczki TL, Griffin MDW. Structural Understanding of Interleukin 6 Family Cytokine Signaling and Targeted Therapies: Focus on Interleukin 11. Front Immunol. 2020;11(July):1–25. doi:10.3389/fimmu.2020.01424

27. Giraldez MD, Carneros D, Garbers C, Rose-John S, Bustos M. New insights into IL-6 family cytokines in metabolism, hepatology and gastroenterology. Nat Rev Gastroenterol Hepatol. 2021;18(11):787–803. doi:10.1038/s41575-021-00473-x

28. Rose-John S. Interleukin-6 family cytokines. Cold Spring Harb Perspect Biol. 2018;10(2):1–17. doi:10.1101/cshperspect.a028415

29. Widjaja AA, Chothani SP, Cook SA. Different roles of interleukin 6 and interleukin 11 in the liver: implications for therapy. Hum Vaccines Immunother. 2020;16(10):2357–2362. doi:10.1080/21645515.2020.1761203

30. Widjaja AA, Viswanathan S, Jinrui D, et al. Molecular Dissection of Pro-Fibrotic IL11 Signaling in Cardiac and Pulmonary Fibroblasts. Front Mol Biosci. 2021;8(September):1–14. doi:10.3389/fmolb.2021.740650

31. Ng B, Dong J, D’Agostino G, et al. Interleukin-11 is a therapeutic target in idiopathic pulmonary fibrosis. Sci Transl Med. 2019;11(511). doi:10.1126/scitranslmed.aaw1237

32. Elias JA, Zheng T, Einarsson O, et al. Epithelial interleukin-11. Regulation by cytokines, respiratory syncytial virus, and retinoic acid. J Biol Chem. 1994;269(35):22261–22268. doi:10.1016/s0021-9258(17)31785-4

33. Corne JM, Holgate ST. Mechanisms of virus induced exacerbations of asthma. Thorax. 1997;52(4):380–389. doi:10.1136/thx.52.4.380

34. Einarsson O, Geba GP, Zhu Z, Landry M, Elias JA. Interleukin-11: Stimulation in vivo and in vitro by respiratory viruses and induction of airways hyperresponsiveness. J Clin Invest. 1996;97(4):915–924. doi:10.1172/JCI118514

35. Ng B, Cook SA, Schafer S. Interleukin-11 signaling underlies fibrosis, parenchymal dysfunction, and chronic inflammation of the airway. Exp Mol Med. 2020;52(12):1871–1878. doi:10.1038/s12276-020-00531-5

36. Fung KY, Louis C, Metcalfe RD, et al. Emerging roles for IL-11 in inflammatory diseases. Cytokine. 2022;149:155750. doi:10.1016/j.cyto.2021.155750

37. She YX, Yu QY, Tang XX. Role of interleukins in the pathogenesis of pulmonary fibrosis. Cell Death Discov. 2021;7(1). doi:10.1038/s41420-021-00437-9

38. Lim WW, Corden B, Ng B, et al. Interleukin-11 is important for vascular smooth muscle phenotypic switching and aortic inflammation, fibrosis and remodeling in mouse models. Sci Rep. 2020;10(1):1–18. doi:10.1038/s41598-020-74944-7

39. Corden B, Adami E, Sweeney M, Schafer S, Cook SA. IL-11 in cardiac and renal fibrosis: Late to the party but a central player. Br J Pharmacol. 2020;177(8):1695–1708. doi:10.1111/bph.15013

40. Katoh M, Katoh M. STAT3-induced WNT5A signaling loop in embryonic stem cells, adult normal rissues, chronic persistent inflammation, rheumatoid arthritis and cancer. Int J Mol Med. 2007;19:273–278.

41. Lopez-Bergami P, Barbero G. The emerging role of Wnt5a in the promotion of a pro-inflammatory and immunosuppressive tumor microenvironment. Cancer Metastasis Rev. 2020;39(3):933–952. doi:10.1007/s10555-020-09878-7

42. Akhmetshina A, Palumbo K, Dees C, et al. Activation of canonical Wnt signalling is required for TGF-β-mediated fibrosis. Nat Commun. 2012;3. doi:10.1038/ncomms1734

43. Beljaars L, Daliri S, Dijkhuizen C, Poelstra K, Gosens R. WNT-5A regulates TGF-β-related activities in liver fibrosis. Am J Physiol - Gastrointest Liver Physiol. 2017;312(3):G219–G227. doi:10.1152/ajpgi.00160.2016

44. Działo E, Tkacz K, Błyszczuk P. Crosstalk between the TGF-β and WNT signalling pathways during cardiac fibrogenesis. Acta Biochim Pol. (3):341–349. 2018;65, doi:10.18388/abp.2018_263

45. Kumawat K, Menzen MH, Bos STI, et al. Noncanonical WNT-5A signaling regulates TGF-β-Induced extracellular matrix production by airway smooth muscle cells. FASEB J. 27(4):1631–1643. doi:10.1096/fj.12-217539 2013

46. Działo E, Czepiel M, Tkacz K, Siedlar M, Kania G, Błyszczuk P. WNT/β-catenin signaling promotes TGF-β-mediated activation of human cardiac fibroblasts by enhancing IL-11 production. Int J Mol Sci. 2021;22(18). doi:10.3390/ijms221810072

47. Di Gregorio J, Robuffo I, Spalletta S, et al. The Epithelial-to-Mesenchymal Transition as a Possible Therapeutic Target in Fibrotic Disorders. Front Cell Dev Biol. 2020;8(December):1–32. doi:10.3389/fcell.2020.607483

48. Ramasamy S, Subbian S. Critical determinants of cytokine storm and type i interferon response in COVID-19 pathogenesis. Clin Microbiol Rev. 2021;34(3). doi:10.1128/CMR.00299-20

49. Morris G, Bortolasci CC, Puri BK, Marx W, Neil AO. Since January 2020 Elsevier has created a COVID-19 resource centre with free information in English and Mandarin on the novel coronavirus COVID-19. The COVID-19 resource centre is hosted on Elsevier Connect, the company’s public news and information. 2020;(January).

50. Coomes EA, Haghbayan H. Interleukin-6 in Covid-19: A systematic review and meta-analysis. Rev Med Virol. 2020;30(6):1–9. doi:10.1002/rmv.2141

51. Ramanathan K, Antognini D, Combes A, et al. Since January 2020 Elsevier has created a COVID-19 resource centre with free information in English and Mandarin on the novel coronavirus COVID-research that is available on the COVID-19 resource centre - including this for unrestricted research re-use a. 2020;(January):19-21.

52. Ghosh MC, Collins GD, Vandanmagsar B, et al. Activation of Wnt5A signaling is required for CXC chemokine ligand 12-mediated T-cell migration. Blood. 2009;114(7):1366–1373. doi:10.1182/blood-2008-08-175869

53. Qi Y, Xu R. Roles of PLODs in collagen synthesis and cancer progression. Front Cell Dev Biol. 2018;6(JUN):1–8. doi:10.3389/fcell.2018.00066

54. Hennet T. Collagen glycosylation. Curr Opin Struct Biol. 2019;56:131–138. doi:10.1016/j.sbi.2019.01.015

55. Ng B, Dong J, Viswanathan S, et al. Fibroblast-specific IL11 signaling drives chronic inflammation in murine fibrotic lung disease. FASEB J. 2020;34(9):11802–11815. doi:10.1096/fj.202001045RR

56. Li H, Xiao H, Lin L, et al. Drug design targeting protein-protein interactions (PPIs) using multiple ligand simultaneous docking (MLSD) and drug repositioning: Discovery of raloxifene and bazedoxifene as novel inhibitors of IL-6/GP130 interface. J Med Chem. 2014;57(3):632–641. doi:10.1021/jm401144z

57. Wei J, Ma L, Lai YH, et al. Bazedoxifene as a novel GP130 inhibitor for Colon Cancer therapy. J Exp Clin Cancer Res. 2019;38(1):1–13. doi:10.1186/s13046-019-1072-8

58. Thilakasiri P, Huynh J, Poh AR, et al. Repurposing the selective estrogen receptor modulator bazedoxifene to suppress gastrointestinal cancer growth. EMBO Mol Med. 2019;11(4):1–15. doi:10.15252/emmm.201809539

59. Moustakas A, Heldin CH. The regulation of TGFβ signal transduction. Development. 2009;136(22):3699–3714. doi:10.1242/dev.030338

60. Blanco-Melo D, Nilsson-Payant BE, Liu WC, et al. Imbalanced Host Response to SARS-CoV-2 Drives Development of COVID-19. Cell. 2020;181(5):1036–1045.e9. doi:10.1016/j.cell.2020.04.026

61. Wang S, Yao X, Ma S, et al. A single-cell transcriptomic landscape of the lungs of patients with COVID-19. Nat Cell Biol. 2021;23(12):1314–1328. doi:10.1038/s41556-021-00796-6

62. Daniel Wendisch OD. SARS-CoV-2 infection triggers profibrotic macrophage responses and lung fibrosis | Elsevier Enhanced Reader. 2021;(January). https://reader.elsevier.com/reader/sd/pii/S0092867421013830?token=2EAB0CEDC3D05073F0BDEA61DAF89DEE6A840FAD49EA1DE224983B2A3EA85F8918CB4794840102657FD62BAEFBF0F236&originRegion=us-east-1&originCreation=20211201230519

63. Naqvi AAT, Fatima K, Mohammad T, et al. Since January 2020 Elsevier has created a COVID-19 resource centre with free information in English and Mandarin on the novel coronavirus COVID-19. The COVID-19 resource centre is hosted on Elsevier Connect, the company’s public news and information. BBA - Mol Basis Dis. 2020;(January):1-17.

64. Krishnamoorthy S, Swain B, Verma RS, Gunthe SS. SARS-CoV, MERS-CoV, and 2019-nCoV viruses: an overview of origin, evolution, and genetic variations. VirusDisease. 2020;31(4):411–423. doi:10.1007/s13337-020-00632-9

65. Lu G, Zhao W, Rao D, Zhang S, Zhou M, Xu S. Knockdown of long noncoding RNA WNT5A-AS restores the fate of neural stem cells exposed to sevoflurane via inhibiting WNT5A/Ryk-ROS signaling. Biomed Pharmacother. 2019;118(July):109334. doi:10.1016/j.biopha.2019.109334

66. Salmena L, Poliseno L, Tay Y, Kats L, Pandolfi PP. A ceRNA hypothesis: The rosetta stone of a hidden RNA language? Cell. 2011;146(3):353–358. doi:10.1016/j.cell.2011.07.014

67. Theocharis AD, Skandalis SS, Gialeli C, Karamanos NK. Extracellular matrix structure. Adv Drug Deliv Rev. 2016;97:4–27. doi:10.1016/j.addr.2015.11.001

68. Ai X, Shen H, Wang Y, et al. Developing a Diagnostic Model to Predict the Risk of Asthma Based on Ten Macrophage-Related Gene Signatures. Biomed Res Int. 2022;2022. doi:10.1155/2022/3439010

69. Palmqvist P. IL-1 and TNF-Regulate in Gingival Fibroblasts. J Dent Res. 2015;87(6):558–563.

70. Janknech R, Hunter T. Activation of the Sap-1a transcription factor by the c-Jun N-terminal kinase (JNK) mitogen-activated protein kinase. J Biol Chem. 1997;272(7):4219–4224. doi:10.1074/jbc.272.7.4219

71. Jagadish N, Rana R, Selvi R, et al. Characterization of a novel human sperm-associated antigen 9 (SPAG9) having structural homology with c-Jun N-terminal kinase-interacting protein. Biochem J. 2005;389(1):73–82. doi:10.1042/BJ20041577

72. Del Sarto J, Gerlt V, Friedrich ME, et al. Phosphorylation of JIP4 at S730 Presents Antiviral Properties against Influenza A Virus Infection. J Virol. 2021;95(20):1–13. doi:10.1128/jvi.00672-21

73. Rui Gui, Huabin Zheng, Liping Ma, Renyi Liu, Xian Lin, Xianliang Ke, Chang Ye XJ. Sperm-Associated Antigen 9 Promotes In fluenza A Virus-. 2022;1(3):1-15.

74. García-García, Tránsito. et al. Impairment of antiviral immune response and disruption of cellular functions by SARS-CoV-2. iScience. Published online 2022. doi:10.1016/j.isci.2022.105444

75. Anders S, Huber W. Differential expression analysis for sequence count data. Genome Biol. 2010;11(10):R106. doi:10.1186/gb-2010-11-10-r106

76. Love MI, Huber W, Anders S. Moderated estimation of fold change and dispersion for RNA-seq data with DESeq2. Genome Biol. 2014;15(12):1–21. doi:10.1186/s13059-014-0550-8

77. R Foundation for Statistical Computing. R: A language and environment for statistical computing. Published online 2020.

78. Oliveros JC. (2007-2015) Venny. An interactive tool for comparing lists with Venn’s diagrams. http://Bioinfogp.Cnb.Csic.Es/Tools/Venny/Index.html. Published online 2015:2015. https://ci.nii.ac.jp/naid/20001505977

79. Zhou Y, Zhou B, Pache L, et al. Metascape provides a biologist-oriented resource for the analysis of systems-level datasets. Nat Commun. 2019;10(1). doi:10.1038/s41467-019-09234-6

