## Supplementary material for "SARS-CoV-2 accessory proteins involvement in inflammatory and profibrotic processes through IL11 signaling": Table S5_List of Primers

**Table S1.**

**List of primers used for qPCR, related to STAR Methods.**

| **Target gen** | **Forward primer (5'-3')** | **Reverse primer (5'-3')** | **Reference** |
| --- | --- | --- | --- |
| ADAMTS1 | CCTCTGTCTGTGTGCAAGGA | GTGGCTCCAGTTGGAATTGT | This work |
| BCL2 | GACAGGAGAAGCCAACAAGC | ACCGCTACTGCAAAGGAGAA | This work |
| CDH2 | AGAGGGGACAACTGCTCTCA | GGCTTTGCTCATGCTACTCC | This work |
| COL1A1 | ACGAAGACATCCCACCAATCAC | TTGCCGTTGTCGCAGACGCAGAT | This work |
| COL3A1 | TGGATGGTGGTTTTCAGTTT | GAAGCTCGGCTGGAGAGAAGT | This work |
| COL4A1 | TCGTGCTACCATCTCTGCAC | ACCTGGTAAGCCTTGGTCCT | This work |
| COL6A1 | GGAAGGCTTGACATGGAAAA | CAGCAAGTCACAGGGACAGA | This work |
| COL11A1 | TGCAATCCCCTCAAATCTTC | ACTCCTTTAAGCCCGGATGT | This work |
| CXCL1 | GGAGCCGTTGGTCAGAAATA | CCTACTGGGCCTCAAATGAA | This work |
| FN1 | GATGCAGACACAGAGCCAAA | CAGTCCTCAGTGGCAGATCA | This work |
| GAPDH | TGGGTGTGAACCATGAGAAG | TGGCAGTGATGGCATGGAC | This work |
| IL11 | GAAACAGCAGGCTACAAAACCACT | ACCTCTCTCCTTTGACCTGGAGAC | This work |
| IL1β | ACAGATGAAGTGCTCCTTCC | CGGCCTGCCTGAAGCCCTTG | This work |
| JUN | CAGCCCACTGAGAAGTCAAACA | CCACCAATTCCTGCTTTGAGA | This work |
| MMP16 | GAAGCACTCTGGGCTTTTTG | GCAACTAGGGTCTGGAGCAG | This work |
| SERPINE1 (PAI-1) | GGCTGACTTCACGAGTCTTT | CTCTCGTTCACCTCGATCTT | This work |
| SNAI1 | GGTTCTTCTGCGCTACTGCT | TAGGGCTGCTGGAAGGTAAA | This work |
| STAT3 | AGCAGCTCCATCAGCTCTACAGT | CTGGCCGCATATGCCCAATCTTG | This work |
| TGFβ | GCGTGCTAATGGTGGAAACCCA | CCGCTTCTCGGAGCTCTGATGTG | This work |
| WNT5A | TGGACAGTGCTCCACAGATTGAT | AATGCATTCTTGGCATCCTTAAA | This work |
| WNT5A-AS | TTGGGGCCACAGAACAAT | GGGGCGACTTTCACCTATTT | This work |
